# Automatic whale counting in satellite images with deep learning

**DOI:** 10.1101/443671

**Authors:** Emilio Guirado, Siham Tabik, Marga L. Rivas, Domingo Alcaraz-Segura, Francisco Herrera

## Abstract

Despite their interest and threat status, the number of whales in world’s oceans remains highly uncertain. Whales detection is normally carried out from costly sighting surveys, acoustic surveys or through high-resolution orthoimages. Since deep convolutional neural networks (CNNs) achieve great performance in object-recognition in images, here we propose a robust and generalizable CNN-based system for automatically detecting and counting whales from space based on open data and tools. A test of the system on Google Earth images in ten global whale-watching hotspots achieved a performance (F1-measure) of 84% in detecting and 97% in counting 80 whales. Applying this cost-effective method worldwide could facilitate the assessment of whale populations to guide conservation actions. Free and global access to high-resolution imagery for conservation purposes would boost this process.

## Whales importance, threats and uncertainties

Whales, which comprise some of the largest animals that have ever existed, have always thrilled humans (*1*, *2*). Whales had and keep an enormous economic and societal value (*3*). More than 13 million whale-watchers were registered in 2008 across 119 countries, generating a global economic activity of US$ 2.1 billion (*4*). Since whales generally are long-living species at high trophic-levels, they play an essential role to structure marine food webs, and to maintain ecosystem functions and services (*5,6*). In the past, commercial whaling depleted whale populations from 66% to 90% from their original numbers, which subsequently caused alterations in marine biodiversity and functions (*7*). To prevent whales from extinction, the signatories of the International Convention for the Regulation of Whaling limited whale hunting to scientific or aboriginal actions since 1946 (*8*). Even though, to date, there still exist a great uncertainty around the number of whales in the oceans and the viability of their populations (*9*). In late 2017, the Species Red List of the International Union for Conservation of Nature (IUCN) (*10*) reported that 22% of 89 evaluated cetacean species were classified as threatened, whereas almost 50% species could not be evaluated due to the lack of data. Hence, a more accurate estimation of whale distribution and population sizes is essential to warrant cetacean conservation (*11*).

## Classical whale-counting methods

The process of identifying and estimating the number of cetaceans is normally carried out 1) *in situ,* from ships, planes or ground stations (*12*), by using visual surveys, acoustic methods (*13*), or a combination of both (*14*); or 2) e*x situ,* by using satellite tracking (*15*, *16*) or photo-interpretation or classical image classification techniques on aerial or satellite images (*17*-*21*). However, these methods are costly, not robust against scenario changes (e.g., different regions or atmospheric conditions), not generalizable to a massive set of images, and often require handcrafted features (*22*). Indeed, biodiversity conservation would certainly benefit from robust and automatic systems to assess species distributions, abundances and movements from space (*23*, *24*).

## Could deep learning help in whale counting?

Deep learning methods, particularly Convolutional Neural Networks (CNNs), could help in this sense since CNNs are already outperforming humans in visual tasks such as image classification and object detection (*25*). CNN models have the capacity to automatically learn the distinctive features of different object classes from a large number of annotated images to later make correct predictions on new images (*26*). Despite building large training datasets is costly, the learning of CNNs on small datasets can be boosted by data-augmentation, which consists of increasing the volume of the training dataset artificially, and by transfer learning, which consists of starting the learning of the network from a prior knowledge rather from scratch (*27*).

Identifying whales from satellite images using CNNs at a global scale is very challenging for several reasons: 1) comprehensive datasets of whale orthoimages to train CNNs do not exist yet; 2) very high-resolution images are expensive and relatively scarce in the marine environment; 3) whales could potentially be confused with other objects such us boats, rocks, waves, or foam; 4) whale postures or swimming movements captured in a snapshot are quite variable since different parts of whale bodies can be emerged or submerged (e.g. lobtailing, blowing, logging, etc.); and 5) occlusions and noise could occur due to clouds, aerosols, haze, sunglint, or water turbidity.

## Objectives

In this work, we propose a robust and generalizable deep learning system for automatically counting whales from space. For this, we built a two-step CNN-based model, where the first step detects the presence of whales and the second step counts the number of whales in satellite images (Fig. 1). To overcome the above mentioned challenges, 1) we combined several open datasets to build an annotated training database of high quality orthoimages of whales and of objects that could be confused with whales, 2) we used data augmentation and transfer learning techniques to make the CNNs robust to image variability, 3) we assessed the effect of whale posture and location on model performance, and 4) we applied the model to free Google Earth coastal imagery in 10 whale watching hotspots as a proof of concept.

**Fig. 1.**
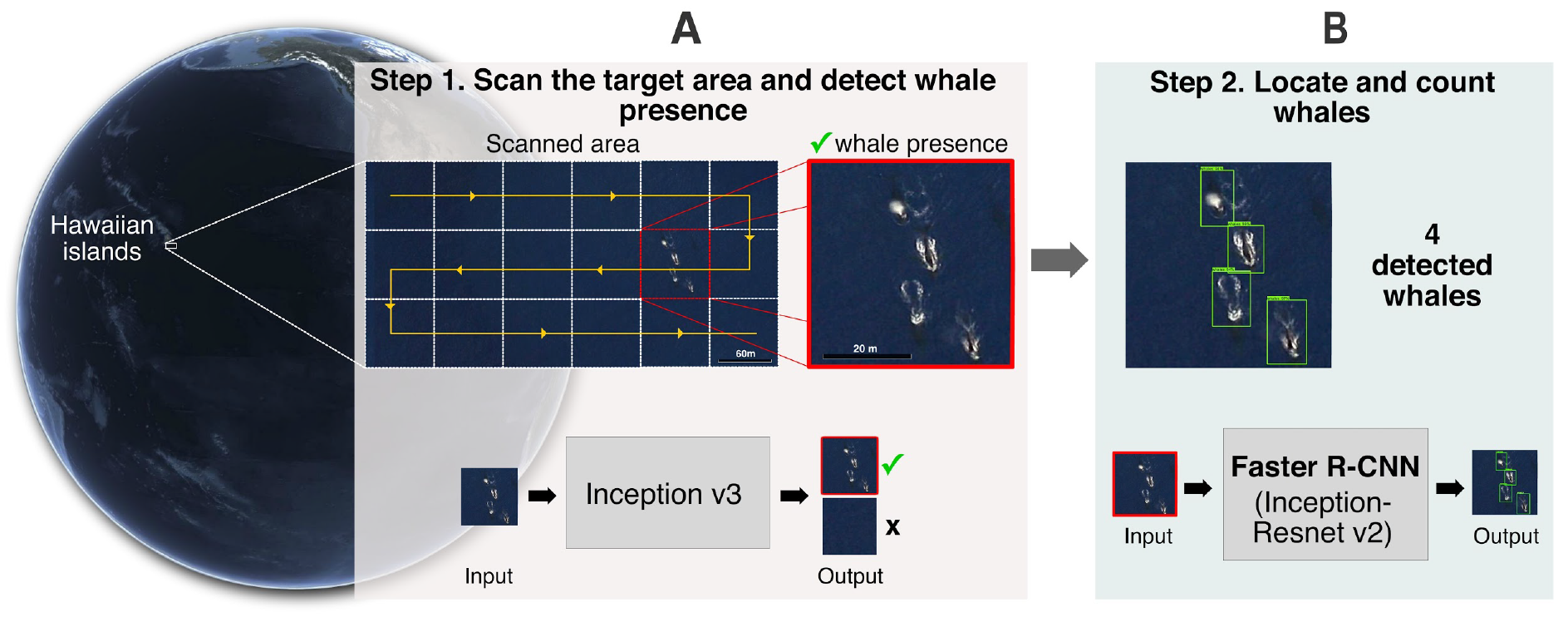
The proposed automatic whale-counting procedure with a two-step CNN-based model. (**A**) The first-step CNN scans the sample area (following the yellow line) to search for the presence of whales in each grid cell (white squares). Only grid cells where the first-step CNN gives high probability for whale presence (red square) are analyzed by (**B**) the second-step CNN, which finally locates and counts individuals (green bounding boxes).

In brief, the first step CNN-based model was applied on a 71 m x 71 m sliding windows-twice the size of blue whales (30 m)- in 10 whale hotspots around the world and outputs the probability of having detected whales in each window (Fig. 1A). The second step CNN-based model analyzed only those windows with high probability of whale presence, localized each whale within a bounding box, and output the number of counted individuals (Fig. 1B). To facilitate its use and to support whale conservation, the CNN-based model was built using open-source software and can be used on free Google Earth images (subjected to terms of service).

## Methods summary

### Training, testing and validating datasets

Since there not exist accessible datasets of orthoimages for whales yet, we had to build two datasets for training the CNN-based models to respectively detect the presence of whales and count their number, and a third dataset for testing and validating the whole procedure. The three datasets were built by combining, preprocessing and labeling images from several sources: Google Earth (*28*), free Arkive (*29*), NOAA Photo Library (*30*), and NWPU-RESISC45 dataset (*31*). For the first step, the training dataset had 2,800 images of the following three object classes (700 images per class): 1) whales, 2) ships, and 3) “water + submerged rocks” (Data S1). For the second step, the same training dataset with 700 images containing whales was used, but each whale was annotated within a bounding box (the total number of bounding boxes was 945).

The dataset for testing and validating the whole procedure consists of RGB satellite images downloaded from Google Earth in 14,148 cells of 71 m x 71 m distributed worldwide. For ships, we selected 400 images from 100 seaports around the world (Fig. 2A, Data S2). For “water + submerged rocks” class, we selected 400 coastal images randomly around the world (Fig. 2A, Data S3). Finally, for whales (Table 1), we downloaded 13,348 cells (Data S4) of 71 m x 71 m from 10 areas that had very high-resolution images at zoom 18 and that are known for marine mammal diversity or whale watching. These areas have been highlighted either as global marine biodiversity hotspots (*32*), marine mammal hotspots (*33*), or irremplazable or priority conservation sites *(11*), and are included within or next to a marine protected area (*34*) (Table 1). Two authors visually inspected all the images to annotate each cell with the name of the corresponding class and with the number of whales. From the 13,348 cells in the 10 hotspots for whale watching, the authors’ visual photo-interpretation revealed whales only in 70 cells.

**Fig. 2.**
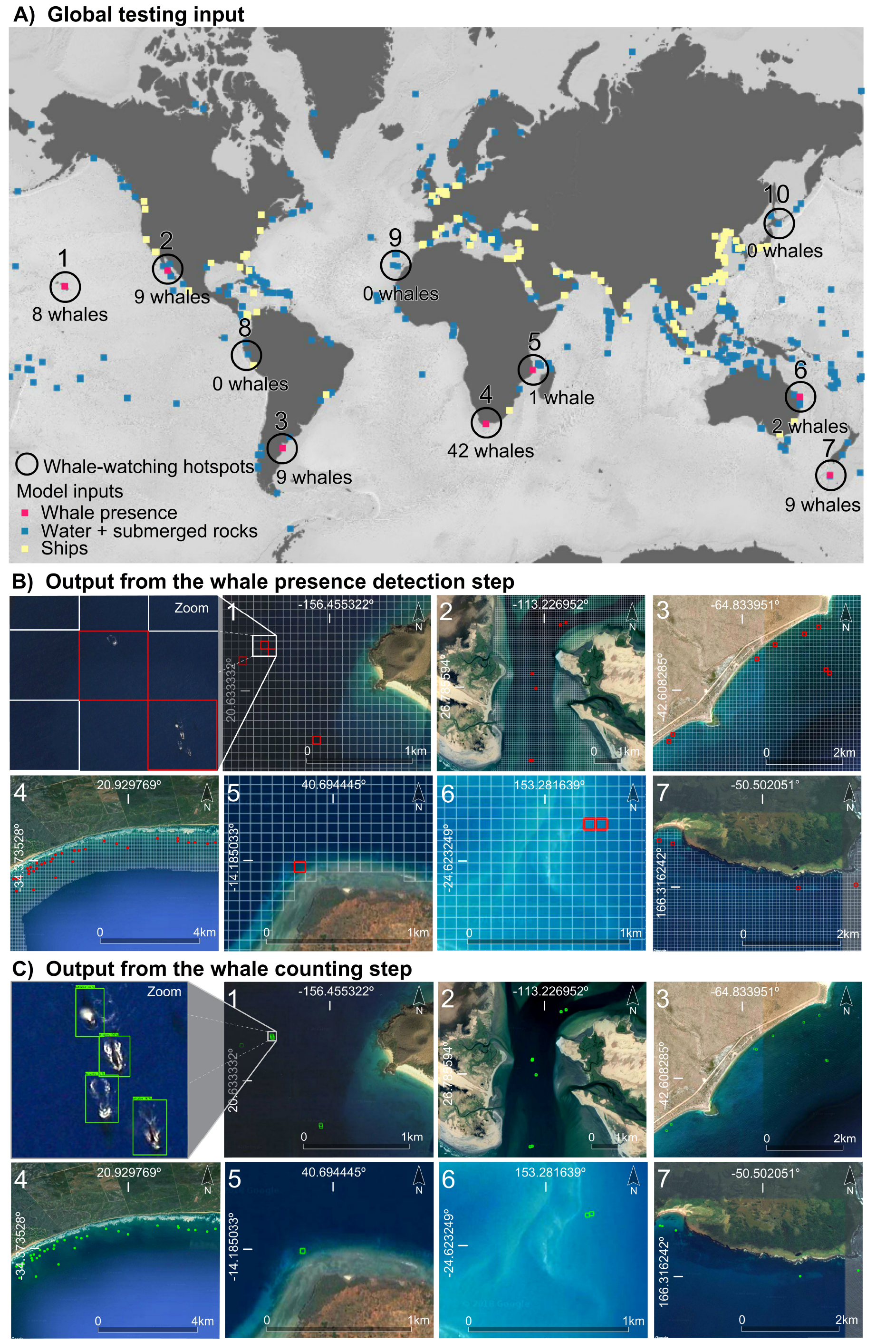
Study sites and illustration of the results from the whale detection and counting models. (**A**) Distribution of 10 marine mammal hotspots (details in Table 1) for whale watching. Red, blue, and yellow cells indicate whale presence, water + submerged rocks, and ships, respectively. (**B**) Detail of the 71 m x 71 m grid cells assessed in coastal areas worldwide. The first step CNN-based model detected the presence of whales in seven of the 10 candidate hotspots (in the three remaining hotspots, high resolution images were not available for the whale watching months). (**C**) Output from the whale counting (Step 2). Detail of the outputs from the second step CNN-based model that located and counted the number of whales (green bounding boxes) in the grid cells where the first-step CNN gave high probability for whale presence.

**Table 1.**
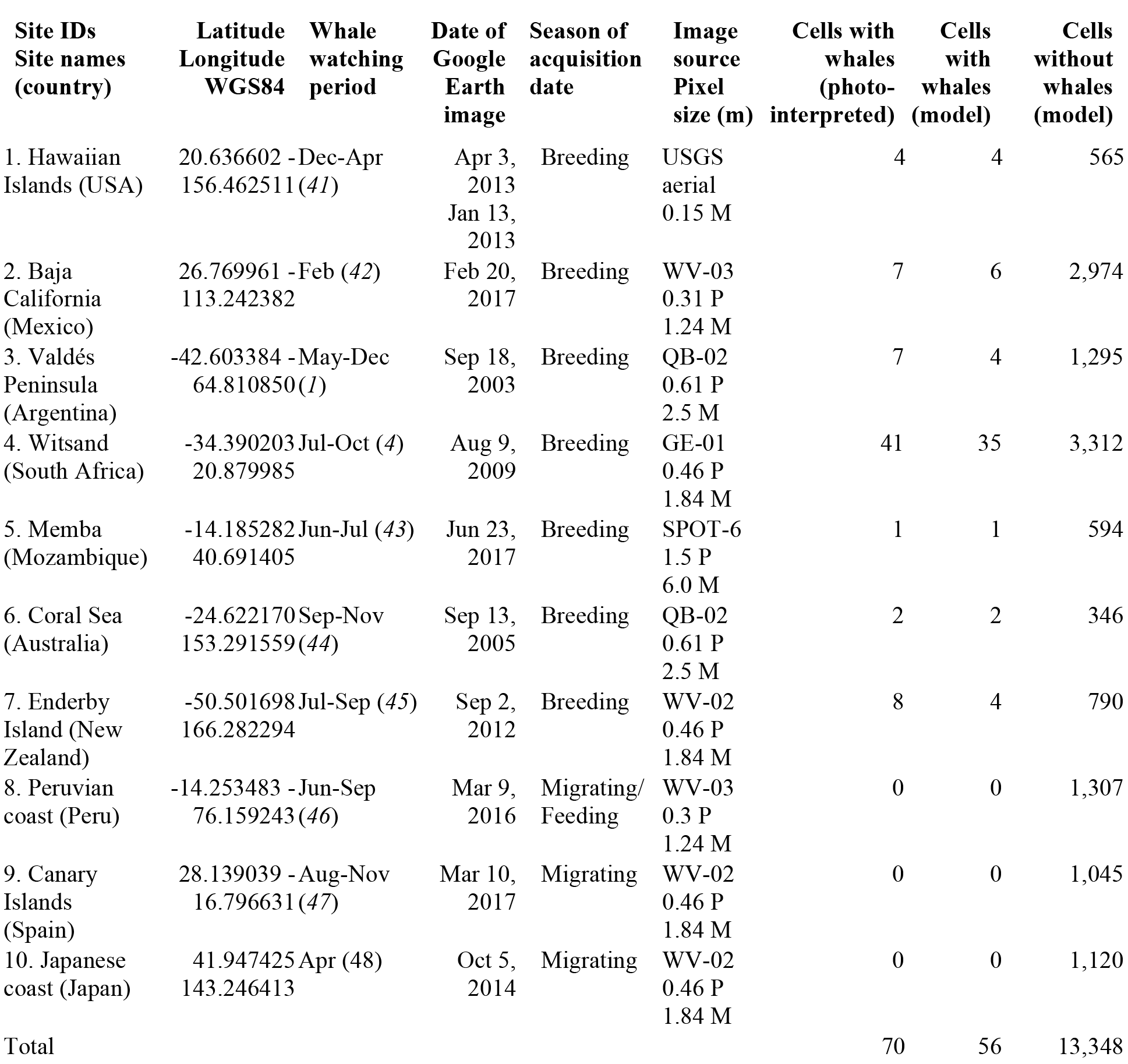
Summary of the inputs and outputs of the whale presence detection model (step 1). Location of the ten sites evaluated. Match between the known whale watching period from the literature and the acquisition date and season of the satellite images in Google Earth. Number of 71 m x 71 m grid cells where the model and the authors detected whale presence and absence. For each image source in Google Earth these metadata are provided: the satellite (GE-01: GeoEye-01; QB-02: QuickBird-2; SPOT-6; USGS: United States Geological Survey orthoimagery; WV-02: WorldView-2; and WV-03: WorldView-3), the pixel size at nadir in meters, and the sensor spectral resolution (M: Multispectral; P: Panchromatic).

### Metrics used in the performance assessment

To evaluate the performance of both CNN-based models, we used these metrics (*35*): positive predictive value, sensitivity, and F1-measure.

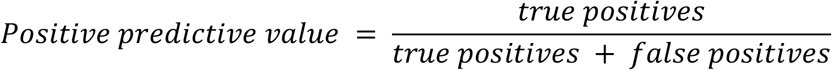

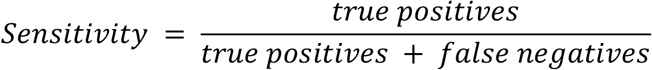

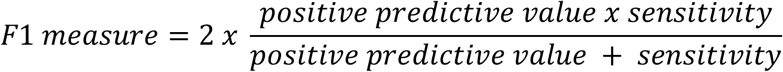

True positives correspond to images that were correctly classified or counted as whales by the models, false positives correspond to images that were classified or counted as whales by the models but actually corresponded to another class, and false negatives correspond to undetected images with whales. In simple terms, high positive predictive value means that the model returned substantially more actual whales than false ones, while high sensitivity means that the model returned most of the actual whales. F1-measure provides a balance between precision and sensitivity.

### Step 1: Whale presence detection phase

When seen from space, whales are often confused with other object classes such as ships and wave foam around partially or entirely submerged rocks. To give the first step CNN-based model the capacity to distinguish between these objects, we addressed the problem as a three-class image classification task. The first model was built using the last version of GoogleNet Inception v3 CNN architecture (*36*), pretrained on the massive ImageNet database (around 1.28 million images, organized into 1,000 object categories). We only removed the two last learnable layers from the network and re-trained them with our dataset, fine-tuning the parameters. To assess whether whale posture, season, and location affected whale presence detection in satellite images, we compared the F1-measure metric across different seasons and locations of the world, and across multiple active and resting swimming movements (*37*), i.e., peduncle, lobtailing, breaching, blowing, spyhopping, logging, and submerged (*38*).

### Step 2: Whale counting phase

We built the second CNN-based model that counts whales by reformulating the problem into an object detection task. We used the detection model Faster R-CNN based on Inception-Resnet v2 CNN architecture (*25*, *39*), pre-trained on the well known COCO (Common Objects in Context) detection dataset, which contains more than 200,000 images organized into 80 object categories (*40*).

## Results

### Whale presence detection and step-1 model validation

The first step CNN-based model that detects the presence of whales reached an average F1-measure of 84% for whales, and of 96% and 97% for the background classes water + submerged rocks and ships, respectively (Table S2, Fig. 3). Only 21.12% of test grid cells containing whales were misclassified as water (18.30 %) or ships (2.82 %). A very small number of water + submerged rocks and ship images were misclassified as whales (1.00 % and 2.25 %, respectively; see Fig. 3).

**Fig. 3.**
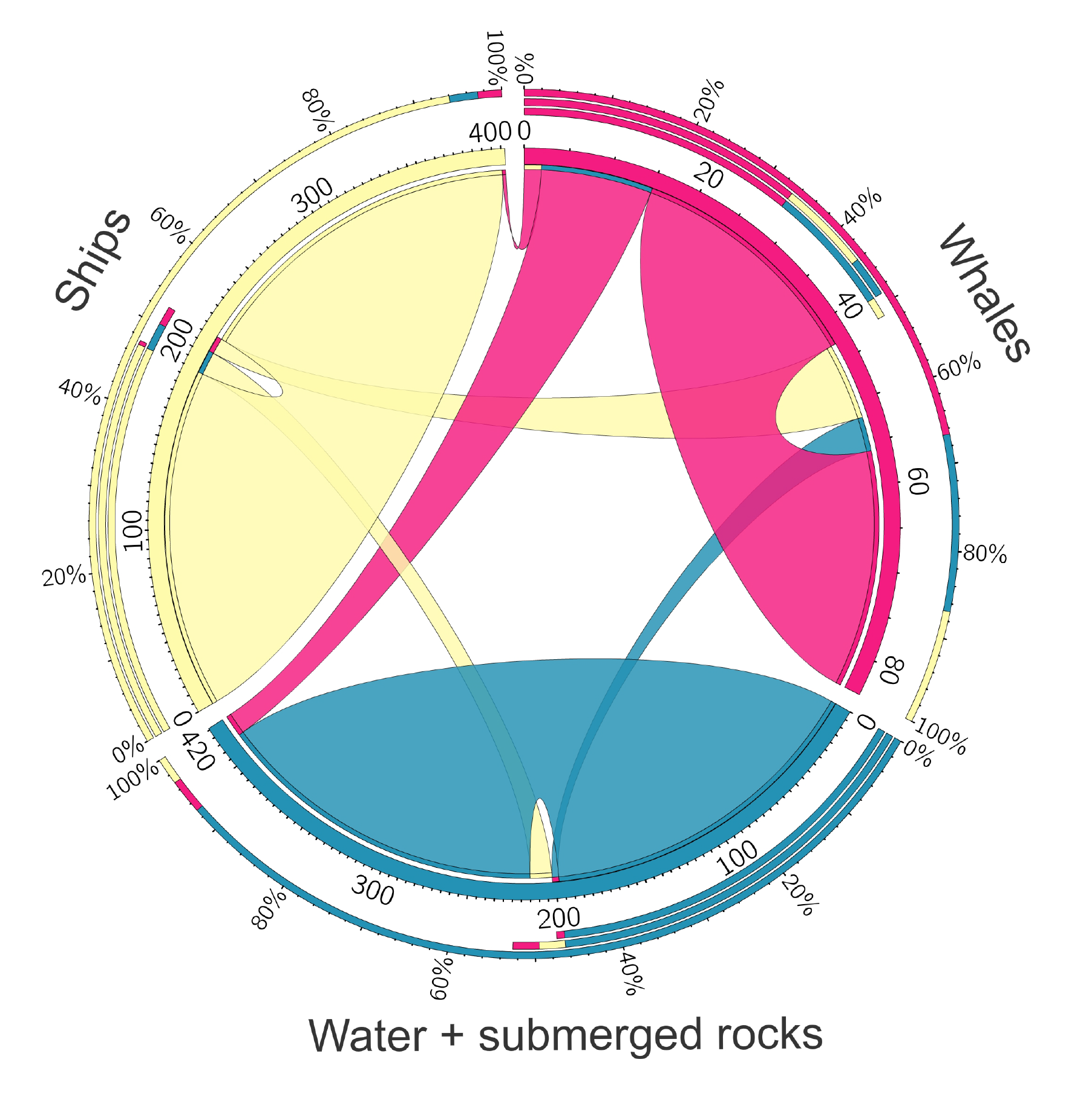
Visualization (Circos plot) of the confusion matrix between the photo-interpreted ground truth and the predictions made by the first step CNN-based model for detecting the presence of whales (in red), ships (yellow), and water + submerged rocks (blue). Only 13 and 1 whale images were classified as water + submerged rocks and as ships, respectively, while only 9 ship images and 4 water + submerged rocks images were classified as whales.

The first step CNN-based model confirmed the presence of whales in seven of the ten whale watching hotspots assessed (see Data S4, Table 1, Fig. 2). The acquisition date of the satellite images available in Google Earth for these seven sites matched the known whale watching period from the literature (*41-48*) (Table 1). In the three sites where the model did not find whales (Peru, Canary Islands, and Japan), the acquisition date of the Google Earth image was not within the known whale watching period but during the migration season. In the Peruvian coast and in the Canary Islands the detection was particularly challenging since the images presented rough sea. However, the author’s photo-interpretation confirmed the absence of whales in the images of Peru, Canary Islands, and Japan (Table 1).

Swimming movements affected the performance of the first step CNN-based model to detect the presence of whales (Fig. 4). Higher detectability (greater than 90% of true positives) was obtained for the following whale postures: blowing, breaching, lobtailing, peduncle, and logging. The lowest detectability occurred for submerged and spyhopping postures (33% and 60% of false negative, respectively; see Fig. 4A). Indeed, the lower performance of the step one model in the Argentinean and New Zealand sites (Table S1, Fig. S1) was due to the much greater frequency of these latter postures in the images (Data S5). Overall, greater number of whales were in the passive swimming movements of logging and submerged (60% of detected whales and 74% of photo-interpreted whales), while the lower number of whales were detected under active movements (Fig. 4A, Data S5).

**Fig. 4.**
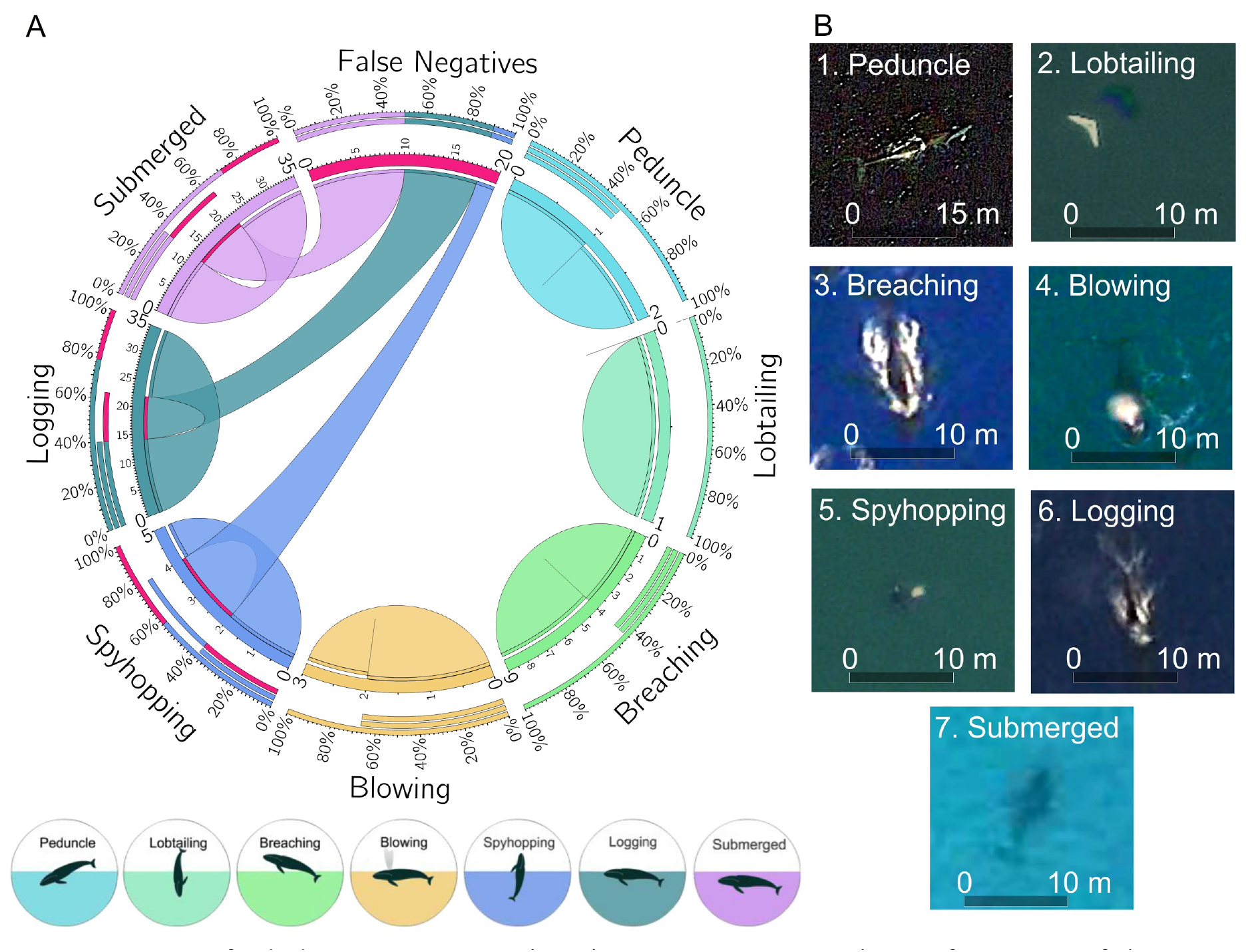
Impact of whale postures or swimming movements on the performance of the Step 1 CNN-based model for detecting the presence of whales. (**A**) The Circos plot shows the distribution of the false negatives (undetected whales, in red color) and true positives across whale postures. Whales under blowing, breaching, lobtailing, and peduncle postures were better detected than under logging, submerged, and spyhopping. (**B**) Example of images for each swimming movement from the detected hotspots (Fig. 2B) at the finest zoom.

### Whale counting and step-2 model validation

The second step CNN-based model localized and counted whales in all the cells where the first step found whale presence, reaching 97 ± 0.04% of F1 measure (Fig. 2C, Table S1). Less than 2% of whales in these cells were not localized and counted by the second step model (one in South Africa). From a total number of 76 whales photo-interpreted in this study across seven hotspots for whale watching around the world, the two-step CNN-based model automatically localized and counted 62 of them, which gives the model an overall global performance of 84 ± 0.13% of F1 measure (Table S1).

## Discussion and conclusions

This study illustrates how global cetacean conservation could benefit from the operational application of deep learning on very high resolution satellite images. Using a two-step convolutional neural network model trained with a reduced dataset and applied on free Google Earth imagery, we managed to automatically detect and count 62 whales in seven hotspots for whale watching around the world, reaching an overall global F1-measure of 89% (F1 measure of 84% for presence detection and 97% for locating and counting). Our results show how the acquisition date of the satellite image and the swimming movement recorded in the image can influence whale presence detection and counting from space. This robust, transparent and automatic method can have direct and wide implications for whale conservation by assessing whale distributions, abundances, and movement behaviours globally from space.

Our satellite-based assessment can complement and be compared with other aerial, marine, and land observations. The coastal images of Google Earth at zoom 18 that we used correspond to a visual altitude of ~254 m, similar to the aerial surveys for grey whales, and up to ~4 km offshore the coast, the maximum distance for whale visual surveys from land (*49*). In whale assessments, such distances are good enough to get reliable estimates of instantaneous presence and relative population abundances (*50*). As new images become available, our method also enables dynamic updates at low cost, to assess seasonal and interannual changes in population sizes, feeding and breeding areas, migratory routes, and distribution ranges around the world.

### CNNs can outperform humans with good quality images

Several studies show that the performance of CNNs can be equal or even better than humans when the quality of the images is good, for instance, for skin cancer detection (*51*), mastering the game of Go (*52*), or generalizing past experiences to new situations (*53*). In general, the quality of the images determines the accuracy of the classification in CNNs (*54*), learning and performing better on higher resolution images (*55*). However, our results show how CNN-based methods trained on high-quality images (see methods section) can also reach good performance in classification and detection on medium-quality images, such as those available for free in Google Earth. In addition, the CNN-based models are robust (*56*) against the differences in spatial and spectral resolutions, and illumination angles across the different satellite sensors used in Google Earth (*29*).

### Limitations of the CNN-based models

The use of free Google Earth imagery is convenient but it also has limitations since these are RGB images rather than multispectral, only available for few dates that may not be within the known whale presence period, are generally constrained to limited locations along coastal areas (up to ~4 km offshore), and are restricted for massive access. These last three limitations must be overcome together with the use of supercomputing for the worldwide “wall-to-wall” application of this method but do not impede its use for local assessments of whales around the world. Image spatial resolution can also limit the application of this method to detect cetaceans shorter than 5 m long (e.g. pilot whales, dolphins, etc.), which would require pixel sizes smaller than 1 m. For example, in our study, higher resolution images tended to give a higher F1-measure (Table S3), though low contrast between whales and surrounding water tended to decreased performance (e.g. New Zealand) and high contrast to improve it (e.g. Mozambique) (Table S3).

Our results showed that the swimming movement and the image acquisition date can also bias the probability of detecting whales. The spatial pattern of whales under blowing, breaching, lobtailing, and peduncle postures showed better detectability than under logging and submerged, when whale bodies can be confused with submerged rocks and seafloor. However, the greater number of whales (both detected by the model and photo-interpreted) in our study were under passive (logging and submerged) instead of active swimming movements, and in images captured during the breeding season. Therefore, the best time to identify whales might be along the breeding season (Table 1), when whales spend more time in surface and in shallow waters (*57*). The effect of overlapped positions between females and calves on their detectability and counting should be further studied. In contrast, the most difficult time might be during migration and in the feeding season (Table 1), when whales are mainly in spyhopping, peduncle, lobtailing, and deeply submerged postures (*58*), and in areas with low contrast between water and whales, or under high sea surface roughness, sea glint, or bad atmospheric conditions (clouds or aerosols).

### A future for conservation with CNNs applied to remote sensing data

The application of CNNs in remote sensing opens a world of possibilities for biodiversity science and conservation (*37*, *59*). The great performance obtained by the CNN-based models trained on and applied to free high-resolution images opens the possibility to automatically process millions of satellite images around the world from whale hotspots, marine protected areas, whale sanctuaries, or migratory routes. Our procedure requires less time and lower cost than the traditional acoustic surveys from ships or the visual surveys from planes and helicopters. The efficiency of remote sensing methods is particularly relevant to save time and money for long-term whale monitoring in remote places, or under difficult circumstances such as whales trapped inside sea ice in polar regions (*60*). The detection of whales using satellite images was already achieved using classical methods (*19*), but their portability to other regions or dates was strongly limited by the necessity of spectral normalization. However, our CNN-based model is easily transferable to any region or image with different characteristics in color, lighting and atmospheric conditions, background, or size and shape of the target objects, and it requires low human supervision, which speeds up the detection process (*36*).

Further research could increase the performance and variety of species identified by our CNN-model. For instance, the model could be improved by increasing the number of samples and variety of atmospheric and sea conditions in the training datasets, by building hierarchical training datasets with different swimming movements across different species (*61*), by using more spectral bands and temporal information (*62*), and by artificially increasing the spatial resolution of the images through rendering (*63*). In addition, as it is a fast and scalable method, it can even be transferred to very high spatial resolution images (<10 cm) captured by unmanned aerial vehicles (UAVs) for the automatic identification of specific individuals (*64*).

A global operationalization of our satellite-based model for whale detection and counting could greatly complement traditional methods (*13*-*21*) to assist whale conservation, to guide marine spatial planning (*65*), or to assess regional (*11*) and global (*33*) priorities for marine biodiversity protection against global change (*66*). In addition, since deep learning is agnostic to the type of image data used, our method could be adapted to identify and quantify other marine species such as seals and sea lions (*67*), penguins (*68*), etc. To boost this process, free access to satellite data is key (*69*). The compromise with biodiversity conservation from corporations such as Google, Microsoft, Planet, Airbus, or DigitalGlobe (*70*) could be materialized through the systematic release of free high resolution aerial and satellite imagery at least from key sites for marine conservation. Even more, the acquisition of these images in pelagic environments does not directly compete with satellites commercial activity, which is usually focused on terrestrial and coastal areas. Having these images available would also make it possible to organize the development of a global database of orthoimages of cetaceans and many other marine vertebrates that could be used to improve the training of our whale detection and counting model or to develop similar models for other marine organisms. Images of the highest spatial resolution (such as WorldView-3 satellite images with a pixel size of 0.3 m) are particularly appropriate for this purpose. This way, satellite and CNN-based detectors of big marine organisms could serve to produce global characterizations of species populations and traits and of community composition as part of the initiative by the Group on Earth Observations - Biodiversity Observation Network (GEOBON) on satellite remote sensing essential biodiversity variables (*71*).

## Funding

S.T. was supported by the Ramón y Cajal Programme of the Spanish government (RYC-2015-18136). S.T., E.G., and F.H. were supported by the Spanish Ministry of Science under the project TIN2017-89517-P. D. A-S. received support from European LIFE Project ADAPTAMED LIFE14 CCA/ES/000612, and from ERDF and Andalusian Government under the project GLOCHARID. D.A-S. received support from NASA Work Programme on Group on Earth Observations - Biodiversity Observation Network (GEOBON) under grant 80NSSC18K0446, from project ECOPOTENTIAL, funded by European Union Horizon 2020 Research and Innovation Programme under grant agreement No. 641762, and from the Spanish Ministry of Science under project CGL2014-61610-EXP and grant JC2015-00316. M.R. received support from International mobility grant for prestigious researchers by (CEIMAR) International Campus of Excellence of the Sea. This work was partially funded by the project BigDaP-TOOLS - Ayudas Fundación BBVA a Equipos de Investigación Científica 2016.

## Author contributions

E.G. and S.T. designed the experiments, collected the data, implemented the code, ran the models, analyzed the results, and wrote the paper. D.A-S., and M.L.R., provided guidance, contributed to the design of the analyses, analyzed the results, and wrote the paper. F.H. provided guidance, contributed to the design of the study, and wrote the paper.

## Competing interests

The authors declare that no competing interests exist in relation to this manuscript.

## Data and material availability statement

The test and training datasets that support our findings are available from Github archive (https://github.com/EGuirado/CNN-Whales-from-Space). Restrictions apply to the availability of the images of the training and validation data, which were used with permission for the current study, but are not publicly available. Some data may be available from the authors upon reasonable request and under written permission from Google Earth, Arkive, or NOAA Photo Library when applicable. To allow reproducibility, in the supplementary information, we provide the metadata of all the images in the training and testing datasets.

The supplementary material listed below is provided under request via

- Tables S1 to S3
- Fig. S1
- Data S1 to S5

